# The multi-BRCT domain protein DDRM2 is required for homologous recombination in plants

**DOI:** 10.1101/2022.04.08.487320

**Authors:** Lili Wang, Chen Yu, Longhui Hou, Yongchi Huang, Xiaoyu Cui, Shijun Xu, Shunping Yan

## Abstract

DNA double-strand breaks (DSBs) are the most toxic DNA damage for cells. Homologous recombination (HR) is a precise DSB repair mechanism as well as a basis for gene targeting using genome-editing techniques. Despite the importance of HR, the HR mechanism in plants is poorly understood. In a genetic screen for DNA Damage Response Mutants (DDRMs), we find that the Arabidopsis *ddrm2* mutant is hypersensitive to DSB-inducing reagents. DDRM2 encodes a protein with four BRCA1 C-terminal (BRCT) domains and is highly conserved in plants including the earliest land plant linage, bryophytes. The plant-specific transcription factor SOG1 binds to the promoter of *DDRM2* and activates its expression, suggesting that *DDRM2* is a direct target of SOG1. In consistence, the expression of *DDRM2* is induced by DSBs in a SOG1-dependent manner. Epistasis analysis indicates that DDRM2 functions downstream of SOG1. Similar to the *sog1* mutant, the *ddrm2* mutant shows dramatically reduced HR efficiency. Our study suggests that the SOG1-DDRM2 module is required for HR, providing new insights into the HR mechanisms in plants and a potential target for improving the efficiency of gene targeting.

**One Sentence Summary:** A genetic screen in Arabidopsis reveals that the multi-BRCT domain protein DDRM2 is required for homologous recombination and is targeted by the master DNA damage response regulator SOG1.

## Introduction

Genome integrity is crucial for the survival of organisms. But genome is constantly challenged by genotoxic insults arising from both exogenous factors (i.e. UV and ionizing radiation) and endogenous factors (i.e. reactive oxygen species and DNA replication errors), leading to various types of DNA lesions. Among them, DNA double-strand breaks (DSBs) are one of the most cytotoxic forms, which, if unrepaired properly, can result in tumorigenesis, premature aging, or cell death (Ciccia and Elledge, 2010). To deal with this, all organisms have evolved complex but elaborate DSB repair mechanisms. Homologous recombination (HR) and non-homologous end-joining (NHEJ) are the two major DSB repair pathways. NHEJ is error-prone and functions throughout all the cell cycle phases. HR is error-free, mainly occurs in the late S and G2 phases (Her and Bunting, 2018). Accumulating evidence suggests that NHEJ and HR compete to repair DSBs (Willis et al., 2018). HR is also a basis of gene targeting using genome-editing tools such as CRISPR/CAS technology (Chen et al., 2019). Due to the low efficiency of HR, it is still a challenge to perform gene targeting in plants (Wolter and Puchta, 2019). There is a great need to improve HR efficiency so as to improve gene targeting efficiency. Therefore, studying the HR mechanism is of both scientific importance and potential implication.

The HR pathway was well-studied in animals. HR is initiated by the recognition of DSBs by the MRE11/RAD50/NBS1 (MRN) complex, which then recruits the protein kinase Ataxia-telangiectasia mutated (ATM). ATM can phosphorylate H2AX, forming an anchor for DNA damage-checkpoint 1 (MDC1), which is also phosphorylated by ATM. Then, MDC1 recruits the ubiquitin E3 ligase RING finger protein 8 (RNF8) to mediate the polyubiquitination of histone H1, which further recruits RNF168 to DSB sites (Mailand et al., 2007). RNF168 ubiquitinates histone H2A to provide a docking site for RAP80 and BRCA1(Doil et al., 2009). BRCA1 then promotes MRN, CtIP, EXO1, and BLM/DNA2 for DNA end resection, producing 3’ single-stranded DNA (ssDNA) overhang, which is protected by RPA proteins. With the help of BRCA2 and other proteins, RPA is replaced by recombinase RAD51 and forms a RAD51 nucleoprotein filament, which performs homologous DNA searching and strand invasion (San Filippo et al., 2008). Recently, it was reported that 53BP1, RIF1, PTIP, and the Shieldin complex inhibit HR and promote NHEJ through protecting DNA end from long-range resection (Noordermeer et al., 2018; Callen et al., 2020). Plants encode the orthologs of ATM, MRN, H2AX, RPA, RAD51, BRCA1, and BRCA2, but lack many other orthologs including MDC1, RNF8, RNF168, 53BP1, RIF1, and Shieldin components. Therefore, it remains largely unknown how plants control HR in plants.

Suppressor of Gamma Response1 (SOG1) is a plant-specific transcription factor and was considered as the functional homolog of p53, a master DNA damage response (DDR) regulator in animals (Yoshiyama et al., 2014). SOG1 plays a crucial role in the transcriptional regulation of DDR genes involved in cell cycle checkpoints, programmed cell death (PCD), and DNA repair (Bourbousse et al., 2018; Ogita et al., 2018). It was shown that SOG1 is required for HR because the HR efficiency in the *sog1* mutant is dramatically reduced compared to wildtype (WT) Arabidopsis (Takahashi et al., 2019).

In this study, we performed a genetic screen for DNA Damage Response Mutants (DDRMs) in Arabidopsis and found that the *ddrm2* mutant is defective in HR. DDRM2 contains four BRCT domains and its biological function is not characterized previously in plants. DSBs induce the expression of *DDRM2* in a SOG1-dependent manner. Consistently, SOG1 binds to the promoter of *DDRM2* and activates its expression. Our study suggests that the SOG1-DDRM2 module is required for HR in plants.

## Results

### *DDRM2* is required for DSB repair

To identify new regulators of DSB repair, we performed a forward genetic screen for *ddrm* mutants using camptothecin (CPT), a topoisomerase I inhibitor that can cause DSBs. The WT Arabidopsis seeds were mutagenized with ethyl methanesulfonate (EMS) and the M2 seeds were grown vertically on medium containing CPT for 8 days. The plants with shorter or longer roots than WT were considered as *ddrm*s. The *atm* mutant was reported to be hypersensitive to the DSB-inducing reagents (Culligan et al., 2006) and thus was used as a positive control. Here we show one of the *ddrm* mutants, *ddrm2-1*. As shown in Figure 1A and 1B, the root length of the *ddrm2-1* mutant was comparable to that of WT and *atm* in absence of CPT. However, the *ddrm2-1* and *atm* mutants showed much shorter roots than that of WT in the presence of CPT, suggesting that the *ddrm2-1* mutant was hypersensitive to CPT. To test whether the *ddrm2-1* mutant responds to other DNA-damaging reagents, we treated the *ddrm2-1* with another DSB-inducing reagent belomycin (BLM) and replication-blocking reagent hydroxyurea (HU). Previous studies suggested that the *atr* mutant was hypersensitive to HU (Culligan et al., 2004) and was used as a positive control. Similar to the case of CPT treatment, the root length of *ddrm2-1* was much shorter than that of WT upon BLM treatment (Supplemental Figure S1A and S1B), indicating that *ddrm2-1* was also hypersensitive to BLM. However, the root length of the *ddrm2-1* mutant was similar to WT upon HU treatment (Supplemental Figure S1C and S1D), suggesting that DDRM2 is not involved in replication stress response. These results suggest that DDRM2 specifically participates in DSB repair.

**Figure 1.**
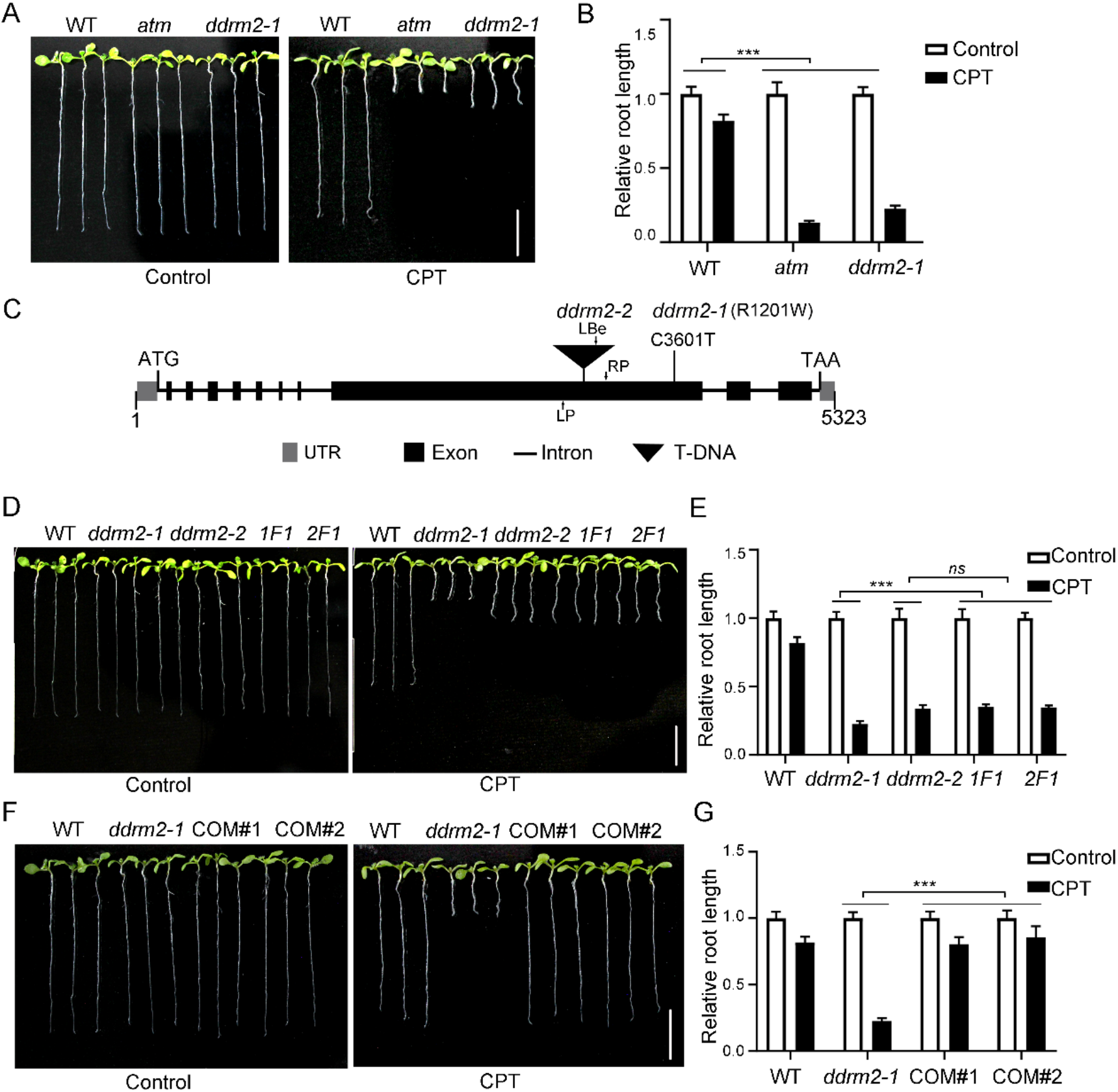
DDRM2 is required for DSB repair. A, D, and F, Pictures of Arabidopsis seedlings grown on 1/2 MS with or without CPT (15 nM) for 8 days. CPT, camptothecin. WT, wildtype. 1F1, the F1 seedlings from a cross of *ddrm2-1* (♂) and *ddrm2-2* (♀). 2F1, the F1 seedlings from a cross of *ddrm2-2* (♂) and *ddrm2-1* (♀). COM, complementation line. Scale bar, 1 cm. B, E, and G, The relative root length of the indicated plants. The relative root length data are represented as means ± SD (n = 10) relative to the values obtained under the control condition. The statistical significance was determined using two-way ANOVA analysis. *\*\*\*, P* < 0.001, *ns*, no significance. All experiments were repeated three times with similar results. C, The genomic structure of *DDRM2*. Black boxes indicate exons and lines indicate introns. ATG and TGA indicate the start and stop codons, respectively. The mutation site of *ddrm2-1*, the T-DNA insertion site of *ddrm2-2*, and the primers used for genotyping are indicated.

To clone *DDRM2*, we used the MutMap strategy (Abe et al., 2012). The *ddrm2-1* mutant was backcrossed to WT. The F2 seedlings were grown vertically on the CPT-containing medium, the plants with shorter roots were sampled as the mutant pool and the plants with longer roots were sampled as the WT pool. The DNA from both the mutant pool and the WT pool was sequenced using the next-generation sequencing technology. The data were analyzed using SIMPLE pipeline (Wachsman et al., 2017), which revealed four candidate genes (Supplemental Table S1). Among them, AT4G02110 encodes a protein with four BRCA1 C-terminal (BRCT) domains. Because many BRCT-domain proteins are involved in DDR (Leung and Glover, 2011), we chose AT4G02110 for further analysis. Firstly, we tested the phenotypes of *ddrm2-2*, another T-DNA insertion mutant of AT4G02110. We found that the *ddrm2-2* mutant was similar to *ddrm2-1* in responses to CPT, BLM, and HU (Figure 1D, 1E, and Supplemental Figure S1A-S1D). Secondly, the reciprocal crosses between *ddrm2-1* and *ddrm2-2* were performed. All the resulting F_1_ seedlings (1F1, 2F1) were hypersensitive to CPT (Figure 1D and 1E), suggesting that *ddrm2-1* and *ddrm2-2* are allelic. Thirdly, we carried out the complementation test by transforming the CDS of *DDRM2* driven by the *CaMV 35S* promoter into the *ddrm2-1* mutant. The resulting transgenic lines (COM) displayed WT-like response to CPT (Figure 1F and 1G), suggesting that *DDRM2* can complement *ddrm2-1*. These results revealed that AT4G02110 is the *DDRM2* gene.

### DDRM2 is a BRCT-containing protein and is induced by DSB-inducing reagents

DDRM2 consists of 1329 amino acids with four BRCT domains: BRCT1 (9-87 aa) and BRCT2 (105-194 aa) at the N-terminus, BRCT3 (1087-1181 aa), and BRCT4 (1209-1296 aa) at the C-terminus. The orthologs of DDRM2 could be identified in all plant species including *Physcomitrium patens* and *Marchantia polymorpha*, two representatives of the earliest land plant linage bryophytes (Figure 2A), suggesting that DDRM2 is a evolutionarily ancient protein. Sequence alignment revealed that the four BRCT domains are highly conserved but the region between BRCT2 and BRCT3 is highly variable. Moreover, the mutation site (R1201W) in *ddrm2-1* is also highly conserved (Supplemental Figure S2). DDRM2 is annotated as a transcription coactivator by TAIR (https://www.arabidopsis.org). However, to our knowledge, its molecular and biological functions have not been characterized.

**Figure 2.**
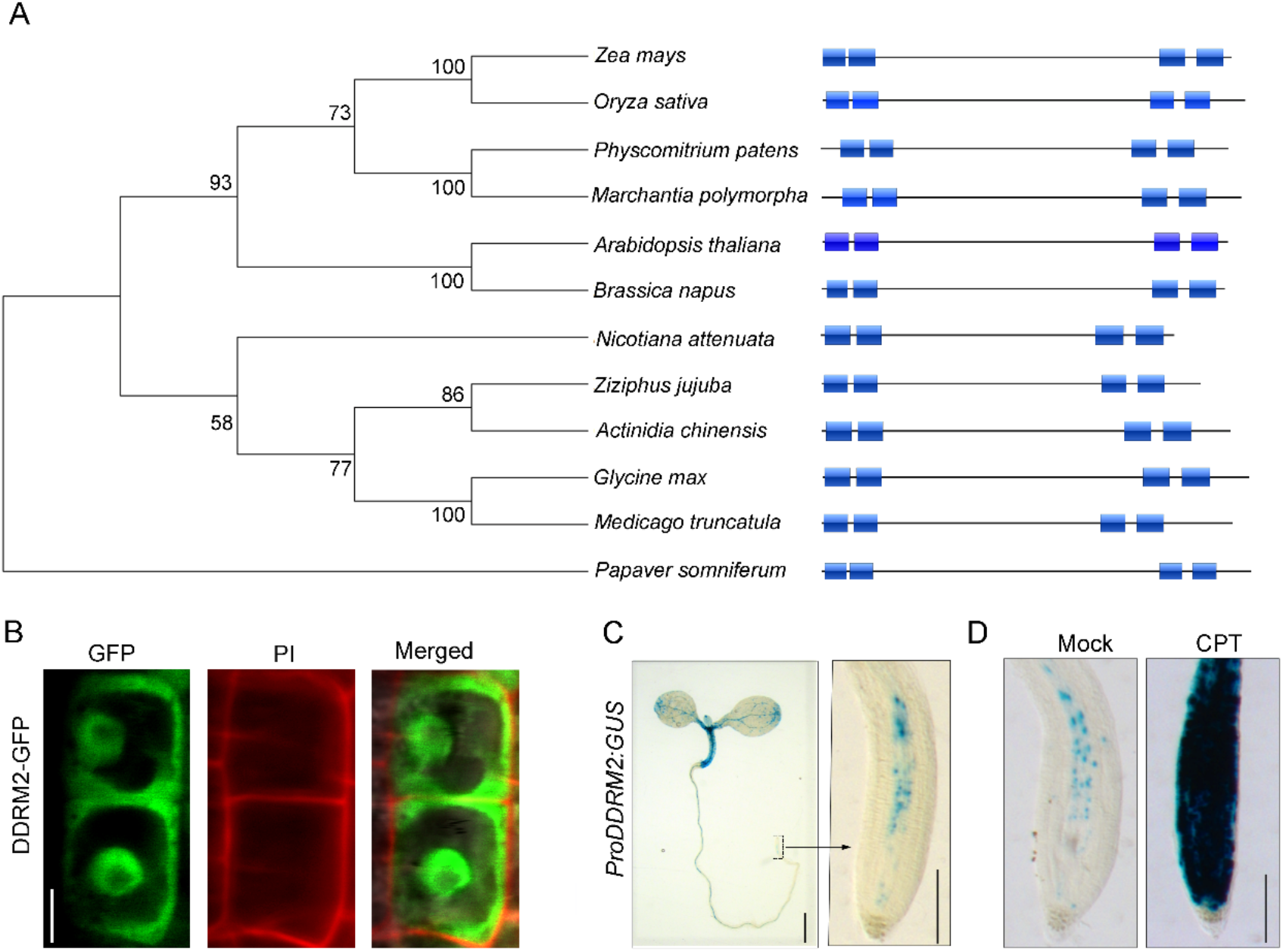
DDRM2 is a conserved nuclear protein and is induced by DNA double-strand breaks. A, Phylogenetic tree of DDRM2 and its orthologs in the indicated species. The BRCT domains are indicated by blue boxes. The amino acid sequences are retrieved from NCBI: *Medicago truncatula* (MTR_1g087290), *Marchantia polymorpha* (MARPO_0010s0152), *Glycine max* (GLYMA_02G013500), Oryza sativa (Os03g0304400), *Nicotiana attenuate* (A4A49_18737), *Actinidia chinensis* (CEY00_Acc15287), *Papaver somniferum* (C5167_015111), *Ziziphus jujuba* (XP_015891972.1), *Zea mays* (NP_001339373.1), *Physcomitrium patens* (XP_024371123.1), *Brassica napus* (XP_013735363.2), and *Arabidopsis thaliana* (AT4G02110). B, Subcellular localization of DDRM2-GFP in the roots of 35S:DDRM2-GFP/*ddrm2-1* transgenic plants. The roots were stained with propidium iodide (PI) to show cell walls. The pictures were captured using confocal microscopy. Scale bar, 5 μm. C, Histochemical staining of transgenic Arabidopsis expressing GUS driven the DDRM2 promoter. Scale bar on the left, 1 mm. Scale bar on the right, 100 μm. D, GUS staining results showing that the expression of *DDRM2* is induced by CPT. The seedlings were treated by 300 nM CPT for 8h. Scale bar, 100 μm.

DDRM2 was predicted to localize in the nucleus by WoLF PSORT (https://wolfpsort.hgc.jp/). To confirm this, the DDRM2-GFP fusion driven by the *CaMV 35S* promoter was transformed into *ddrm2-1*. The resulting transgenic plants were similar to WT upon CPT treatment (Supplemental Figure S3), suggesting that DDRM2-GFP is biologically active. DDRM2-GFP was detected both in the nucleus and cytoplasm through confocal microscopy analysis (Figure 2B).

To test the expression patterns of *DDRM2*, the *GUS* gene driven by *DDRM2* native promoter (*proDDRM2:GUS*) was transformed into WT plants. Among 16 transgenic lines, 15 of them showed a similar expression pattern. As shown in Figure 2C, the expression of *DDRM2* was highest in hypocotyls in the 6-day-old seedlings. It can also express in cotyledons and roots, especially in the vascular tissues. To our surprise, *DDRM2* was weakly expressed in the root meristem, where many genes involved in DNA repair are highly expressed. This result suggested that *DDRM2* may be induced upon DNA damage treatment. Indeed, the expression of *DDRM2* in the root tips was dramatically enhanced after CPT treatment (Figure 2D). This result was confirmed through quantitative reverse transcription-PCR assay (qRT-PCR) (Figure 3A).

**Figure 3.**
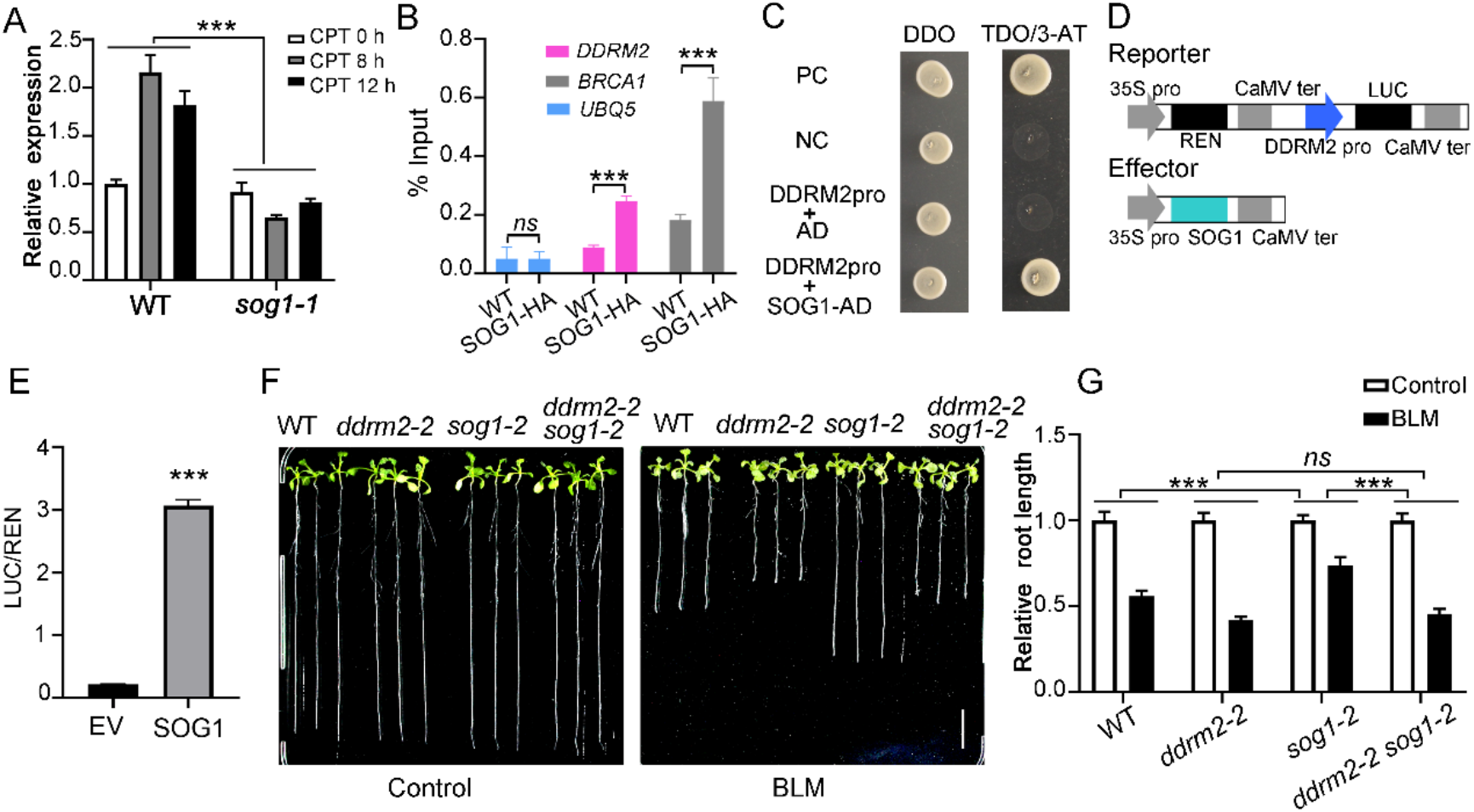
*DDRM2* is a target gene of SOG1. A, The relative expression of *DDRM2* in WT and *sog1-1* seedlings treated with 40 nM CPT for different times (0, 8, 12 h). The relative expression level of *DDRM2* was determined by qRT-PCR analysis using *ubiquitin 5* (*UBQ5*) as an internal standard. B, SOG1 associates with *DDRM2* promoter in ChIP-qPCR assays. The anti-HA antibody was used to perform immunoprecipitation. *UBQ5* was used as a negative control. *BRCA1* was used as a positive control. C, SOG1 binds to the promoter of *DDRM2* in yeast one-hybrid assay. The promoter fragment of *DDRM2* was cloned into *pHis2*.*1* vector (DDRM2pro). SOG1 was cloned into *pGADT7-rec2* vector (SOG1-AD). *pGADT7-Rec-53* and *pHis2*.*1-P53* were used as a positive control (PC). *pGADT7-Rec-53* and *pHis2*.*1* were used as a negative control (NC). DDO, double dropout (SD/-Trp/-Leu) medium; TDO, triple dropout (SD/-Trp/-Leu/-His) medium; 3-AT, 80 mM 3-amino-1,2,4-triazole. D, Schematic representation of the constructs used in dual-luciferase assays. E, SOG1 activates *DDRM2* expression in dual-luciferase assays. The reporters and effectors were co-expressed in Arabidopsis protoplasts, and both REN and LUC activity were measured. The relative LUC activities normalized to the REN activities are shown (LUC/REN). EV, empty vector. Data represent mean ± SE of three biological replicates. F, Pictures of Arabidopsis seedlings grown on 1/2 MS for 5 days and then were transferred onto 1/2 MS with or without BLM (2 μM) for another 6 days. Scale bar, 1 cm. G, The relative root length of the indicated plants. The relative root length data are represented as means ± SD (n = 10) relative to the values obtained under the control condition. The statistical significances in A and G were determined using two-way ANOVA analysis. *\*\*\*, P* < 0.001, *ns*, no significance. The statistical significances in C and E were determined using one-tailed Student’s t-test. *\*\*\*, P* < 0.001, *ns*, no significance. All experiments were repeated three times with similar results.

### *DDRM2* is a target gene of SOG1

It has been shown that the plant-specific transcription factor SOG1 plays a central role in the transcriptional regulation of DNA repair genes (Bourbousse et al., 2018; Ogita et al., 2018). To test whether the DSB-induced expression of *DDRM2* is dependent on SOG1, we examined the DDRM2 expression levels in the *sog1-1* mutant through qRT-PCR analyses. As expected,, we found that the induced expression of *DDRM2* was abolished in the *sog1-1* mutant (Figure 3A). Precious high-throughput studies suggested that DDRM2 is a target gene of SOG1 (Bourbousse et al., 2018; Ogita et al., 2018). To further confirm this, we performed the Chromatin Immunoprecipitation-qPCR (ChIP-qPCR) assays using the transgenic plants expressing SOG1-HA driven by the *CaMV 35S* promoter. The well-known SOG1 target gene *BRCA1* was used as a positive control. Indeed, the promoter regions of both *DDRM2* and *BRCA1* were significantly enriched in the SOG1-HA transgenic plants compared with WT (Figure 3B). To test whether SOG1 binds the promoter of *DDRM2* directly, we performed yeast-one-hybrid (Y1H) assays. The promoter of *DDRM2* (1000 bp upstream of the start codon) was cloned into *pHis2* vector and *SOG1* was cloned into *pGADT7* (AD) vector. As shown in Figure 3C, compared with the negative controls, the yeasts expressing SOG1-AD could grow on the selective medium (TDO+3AT), suggesting that SOG1 can directly bind to the promoter of *DDRM2*. To examine whether SOG1 activates *DDRM2*, the dual-luciferase assays were performed in Arabidopsis protoplasts. The reporter vector contains a firefly luciferase (LUC) gene driven by the *DDRM2* promoter and a renilla luciferase (REN) gene driven by the *CaMV 35S* promoter. The effector vector encodes SOG1 driven by the *CaMV 35S* promoter (Figure 3D). As shown in Figure 3E, the expression ratio of LUC and REN was significantly higher when SOG1 was expressed compared with the empty vector (EV) control. These results strongly suggested that SOG1 can directly bind to the promoter of *DDRM2* and activate its expression, indicating that *DDRM2* is a target gene of SOG1.

The *sog1* mutant was more resistant to BLM than WT (Yoshiyama *et al*, 2017) and *ddrm2* was more sensitive to BLM than WT (Supplemental Figure S1), which allowed us to perform epistasis analysis. We generated the *ddrm2-2 sog1-2* mutant through genetic crossing. As shown in Figure 3F and 3G, the sensitivity of the *ddrm2-2 sog1-2* double mutant to BLM was similar to that of *ddrm2-2*, indicating that DDRM2 functions downstream of SOG1.

### DDRM2 is required for HR

It was shown that the HR efficiency in the *sog1* mutant was reduced compared with WT (Takahashi et al., 2019). Since DDRM2 functions downstream of SOG1, it is likely that DDRM2 is also involved in HR. To test this, we compared the HR efficiency of *ddrm2-2* and WT using a well-established HR reporter system IU.GUS (Roth et al., 2012). The reporter (R) line harbors an *I-SceI* restriction site within the two non-functional β-glucuronidase (GUS) fragments and a nearby donor sequence (U) in inverted orientation. The trigger (T) line expresses the endonuclease *I-SceI*. In the crossed (RxT) line, DSBs are generated at the *I-SceI* site and HR-mediated DSB repair will reconstitute a functional GUS gene, resulting in blue sectors after GUS staining (Figure 4A). As shown in Figure 4B and 4C, the HR efficiency in the *ddrm2-2* mutant was reduced to 40% of that in WT, indicating that DDRM2 is required for HR.

**Figure 4.**
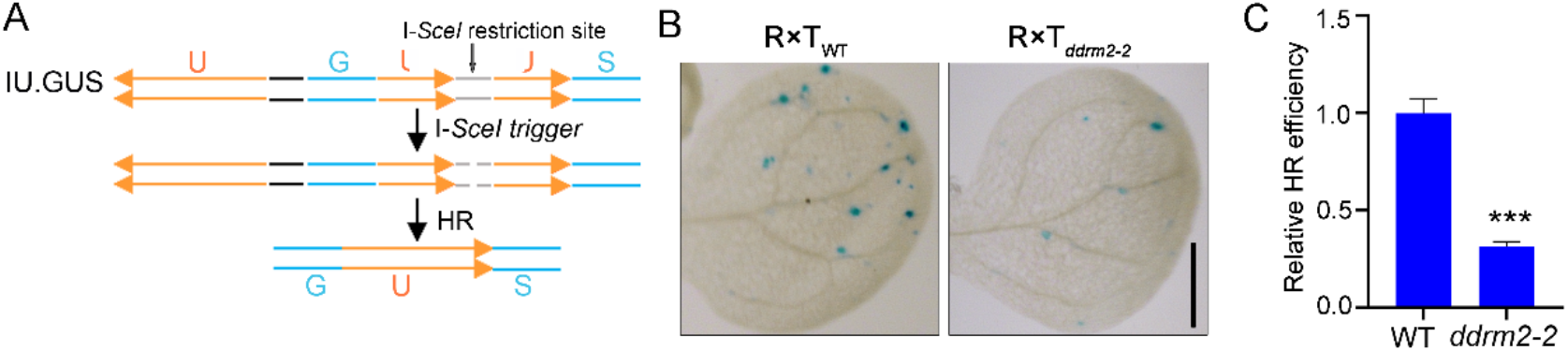
DDRM2 is required for homologous recombination. A, Schematic representation of the IU.GUS reporter system. The reporter (R) line harbors an I-*SceI* restriction site located between two nonfunctional GUS fragments and a nearby donor sequence (U) in inverted orientation. A single DSB is introduced when the reporter line (R) is crossed with the DSB-triggering (T) line that expresses the I-*SceI* endonuclease. When the DSB is repaired through HR, the functional GUS is restored. B, Representative GUS staining images of cotyledons. The reporter line and trigger line in either *ddrm2-2* or WT background were crossed and the F1 seedlings were used for scoring. Scale bar, 1 mm. C, The relative HR efficiency. The HR efficiency in WT was set to 1.0. Data represent mean ± SE of 64 plants in each genetic background. The statistical significances were determined using Student’s t-test. *\*\*\*, P* < 0.001.

## Discussion

Repair of DSBs is critical for cell survival. To date, numerous proteins involved in DSB repair have been identified both in animals and yeasts (Gupta et al., 2018). However, many of them could not be identified in plants (Hu et al., 2016). Therefore, how plants repair DSBs remains elusive. In this study, we found that DDRM2 is an essential regulator of HR. DDRM2 is targeted by the plant-specific protein SOG1 (Figure 3), which is a master regulator of plant DDR. In a recent study, we demonstrated that a plant-specific E3 ubiquitin ligase DDRM1 ubiquitinates and stabilizes SOG1 to regulate HR (Wang et al., 2022). Therefore, DDRM1, SOG1, and DDRM2 function in the same pathway to regulate HR, with DDRM1 upstream of SOG1 and DDRM2 downstream of SOG1. Our study provides not only new insights into HR mechanisms but also potential targets for improving the efficiency of gene targeting.

BRCT domain is originally identified in tumor suppressor BRCA1 and is considered as a protein interaction domain. Recent studies suggest that BRCT domain can not only recognize phosphorylated peptides, but also mediate the interactions with the non-phosphorylated protein, DNA, and poly (ADP-ribose) (Leung and Glover, 2011). In animals and yeasts, BRCT domain is present in many DDR proteins such as 53BP1, MDC1, BRAD1, PARP1, XRCC1, LIG4, TOPBP1, and PTIP (Zhang et al., 2005; Singh et al., 2008). These proteins contain various number of BRCT domains or contain other protein domains. A previous bioinformatics study suggested that DDRM2 and MEI1, which contains five BRCT domains, are two homologs of human HsTOPBP1 (Shultz et al., 2007). HsTOPBP1 contains nine BRCT domains and plays essential roles in DNA replication (Kumagai et al., 2010) and replication stress responses (Bigot et al., 2019). However, the *ddrm2* mutants grow normally and are not sensitive to HU-induced replication stress (Figure 1 and Supplemental Figure S1), suggesting that DDRM2 is divergent from HsTOPBP1.

Although our data clearly showed that DDRM2 is required for HR (Figure 4), how it regulate HR remain to be further elucidated. Given that DDRM2 contain four BRCT domains, it is likely that DDRM2 functions in HR by interacting with other proteins. Therefore, one of the future direction is to identify DDRM2-interacting proteins. Since DDRM2 is highly conserved in plants, it will also be interesting to test whether the DDRM2 homologues from other plant species, especially crops, are essential for HR.

## Materials and Methods

### Plants materials and growth conditions

*Arabidopsis thaliana* mutants used in this study are in the Columbia (Col-0) background. The *ddrm2-2* (SALK_051265), *atm* (SALK_006953), *ku70* (SALK_123114C), *sog1-2* (GK-143A02), *rad5*1 (GABI_134A01), *brca1* (SALK_014731), and *atr* mutant (SALK_032841) mutants were obtained from Arabidopsis Biological Resource Center (ABRC). The *sog1-1* mutant was described previously (Yoshiyama et al., 2009). Seeds were sterilized with 2% PPM (Plant Cell Technology), stratified at 4 °C in the dark for 2 days, and then plated on 1/2 Murashige and Skoog (MS) medium containing 1% (w/v) sucrose and 0.4% (w/v) phytagel. The plants were grown under long-day conditions (16 h of light and 8 h of dark) at 22°C in a growth chamber.

### Genetic screen for *ddrm*

The Col-0 seeds were mutagenized with 0.2% ethyl methanesulfonate (EMS) and grown in soil to produce M2 seeds. M2 seeds were grown vertically on 1/2 MS medium containing 15 nM CPT for 8 days. The plants with longer or shorter roots than WT were considered to be *ddrms*.

### Cloning of *DDRM2*

To clone *DDRM2*, the *ddrm2-1* mutant was crossed with WT. The F2 seedlings were grown vertically on the CPT-containing medium. The plants with shorter root were considered as the mutant pool and the plants with longer roots were considered as the WT pool. These plants were transferred into the soil and grown for two weeks. The leaf discs were separately pooled for genomic DNA extraction. The DNA from both WT and mutant pools were subjected to next-generation sequencing (NGS) by Novogene. The sequence data were analyzed using SIMPLE pipeline to obtain candidate genes (Wachsman et al., 2017).

### Phylogenetic tree analyses

Alignment of protein sequences was performed with ClustalX (gap open penalty: 10; gap extension penalty: 0.1; protein weight matrix: BLOSUM). The phylogenetic analysis was performed with MEGA6 (Tamura et al., 2013). Evolutionary relationships were deduced using the Neighbour-Joining method with bootstrap values (1,000 replicates). Evolutionary distances were computed with the Jones-Taylor-Thornton model with default values (rate among sites: uniform rates; gap/missing data treatment: complete deletion; ML heuristic method: NNI).

### Plasmid Construction

All the vectors used in this study were constructed using a Lighting Cloning system (Biodragon Immunotechnology, China). For complementation assay, CDS of *DDRM2* was amplified and cloned into *pCAMBIA1301* vector. For subcellular localization assay, CDS of *DDRM2* was cloned into modified *pFGC5941*vector with a GFP tag under the control of *CaMV 35S* promoter. For GUS staining, the *DDRM2* promoter (2,000 bp upstream of start codon ATG) was cloned into *pCAMBIA2300-YG* vector. Primers used for the construction of vectors were listed in Supplemental Table S2.

### Subcellular localization

The roots of 35S: DDRM2-GFP/*ddrm2-1* transgenic seedlings were stained with propidium iodide (PI), and the PI and GFP fluorescence signals were observed using confocal microscopy (Leica TCS SP8, Germany).

### GUS staining

The seedlings were incubated in GUS staining solution (100 mmol/L sodium phosphate buffer pH 7.0, 0.1% Triton X-100, 0.5 mmol/L potassium ferrocyanide, 0.5 mmol/L potassium ferricyanide, and 0.5 mmol/L X-Gluc) in the dark at 37°Cfor 12 hours. After that, the samples were washed several times with 70% ethanol and observed under a light microscope (Nikon SMZ1000, Japan).

### Quantitative RT-PCR

Total RNA was extracted using TRIzol Reagent (Invitrogen, USA). The reverse-transcription reaction was performed using HiScript III 1st Strand cDNA Synthesis Kit (+gDNA wiper) according to the manufacturer’s protocol (Vazyme, China). The quantitative PCR assays were performed on the CFX ConnecTM Real-Time PCR Detection System (Bio-Rad, USA) using ChamQ Universal SYBR qPCR Master Mix (Vazyme, China). The *ubiquitin 5* (*UBQ5*) was used as a reference gene. The relative expression level was calculated by the 2^-ΔΔCT^ method.

### Yeast one-hybrid

Yeast one-hybrid assays were performed using the Matchmaker Yeast One-Hybrid System (Clontech). The DNA fragment 1,000 bp upstream of the start codon ATG of *DDRM2* was cloned into *pHis2*.*1* vector and SOG1 was cloned into the pGADT7 vector. They were co-transformed into the yeast strain Y187. Growth performances of the transformants on the SD/-Leu/-Trp and SD/-Leu/-Trp/-His media containing 3-aminotriazole (3-AT) were observed to evaluate the DNA– protein interactions.

### ChIP-qPCR

ChIP assays were performed as described previously with some modifications (Zhao et al., 2020). Briefly, 0.5 g of 10-day-old *35S:SOG1-HA* transgenic seedlings grown on MS plates were cross-linked under vacuum with 1% formaldehyde for 15 min at room temperature. The crosslinking reaction was stopped by adding glycine to a final concentration of 0.2 M. The seedlings were washed with ice-cold water and then ground in liquid nitrogen. The fine powder was lysed in 750 μl of Buffer S (50 mM HEPES-KOH pH7.5, 150 mM NaCl, 1 mM EDTA, 1% Triton X-100, 0.1% sodium deoxycholate, 1% SDS) for 10 min at 4°C. The homogenate was mixed with 3.75 ml of Buffer F (50 mM HEPES-KOH pH7.5, 150 mM NaCl, 1 mM EDTA, 1% Triton X-100, 0.1% sodium deoxycholate), and then the mixture was sonicated to shear DNA into 250 to 600 bp fragments using Bioruptor Plus sonication system (Diagenode, Belgium). The lysates were centrifuged at 20,000 × g for 10 min at 4°C, and the supernatant was transferred to a new tube containing protein G beads. Then the precleared chromatin was incubated with the anti-HA antibody/protein G beads complexes overnight at 4°C. The immunoprecipitated chromatin was washed subsequently with low-salt ChIP buffer (50 mM HEPES-KOH, 150 mM NaCl, 1 mM EDTA, 1% Triton X-100, 0.1% sodium deoxycholate, 0.1% SDS), high-salt ChIP buffer (low-salt ChIP buffer replaced 150 mM NaCl with 350 mM NaCl), ChIP wash buffer (10 mM Tris-HCl pH 8.0, 250mM LiCl, 0.5% NP-40, 1 mM EDTA, 0.1% sodium deoxycholate), and TE buffer. The protein-DNA complexes were eluted from beads by adding 100 μl of freshly prepared ChIP Elution buffer for 15 min at 65 °C. The purified DNA samples were subjected to qPCR analysis.

### Dual-luciferase assay

The dual-luciferase assays were performed as described previously (Wang et al., 2018). The DNA fragment 1000 bp upstream of the start code ATG of *DDRM2* was cloned into *pGreenII 0800-LUC* reporter vectors as a transcriptional fusion with the firefly luciferase (LUC). In the same vector, the renilla luciferase (REN) reporter gene driven by the *CaMV 35S* promoter was used as an internal standard in each transformation. SOG1 was cloned to the effector vector *pFGC5941* driven by the *CaMV 35S* promoter. The reporter and effector constructs were co-expressed in Arabidopsis protoplasts. The luciferase activities were measured on the Mithras LB 940 multimode microplate reader (BERTHOLD technologies, Germany) using Dual-Luciferase Reporter Assay System (Promega, USA).

### HR efficiency assay

The HR efficiency assay was performed as described previously (Roth et al., 2012). The IU.GUS reporter (R) line and the DSB-triggering (T) line were introduced into *ddrm2-2* background. The progenies of RxT crossed plants were then used for GUS staining.

## Supplemental data

**Supplemental Figure S1**. The *ddrm2* mutants are hypersensitive to BLM, but not HU (Supports Figure 1).

**Supplemental Figure S2**. The sequence alignment of DDRM2 and its orthologs in other plant species (Supports Figure 2).

**Supplemental Figure S3**. DDRM2-GFP fusion protein driven by the *CaMV 35S* promoter can rescue *ddrm2-1* (Supports Figure 2).

**Supplemental Table S1**. Candidate genes revealed by SIMPLE analysis.

**Supplemental Table S2**. Primers used in this study.

## Acknowledgments

We thank Prof. Yijun Qi for gifting the HR reporter system. The work was supported by the National Natural Science Foundation of China (32070312, 31970311, 31800216), Thousand Talents Plan of China-Young Professionals, Huazhong Agricultural University Scientific & Technological Self-innovation Foundation (2014RC004).

## Author contributions

S.Y. designed the research, L.W., C.Y., L.H., Y.H., X.C., and S.X. performed experiments and analyzed data, L.W. and S.Y. wrote the manuscript.

## Notes

### Competing Interest Statement

The authors have declared no competing interest.

