## Supplemental Figure S1-3, Supplemental Table S1-2 for "The multi-BRCT domain protein DDRM2 is required for homologous recombination in plants"

### Supplementary Data

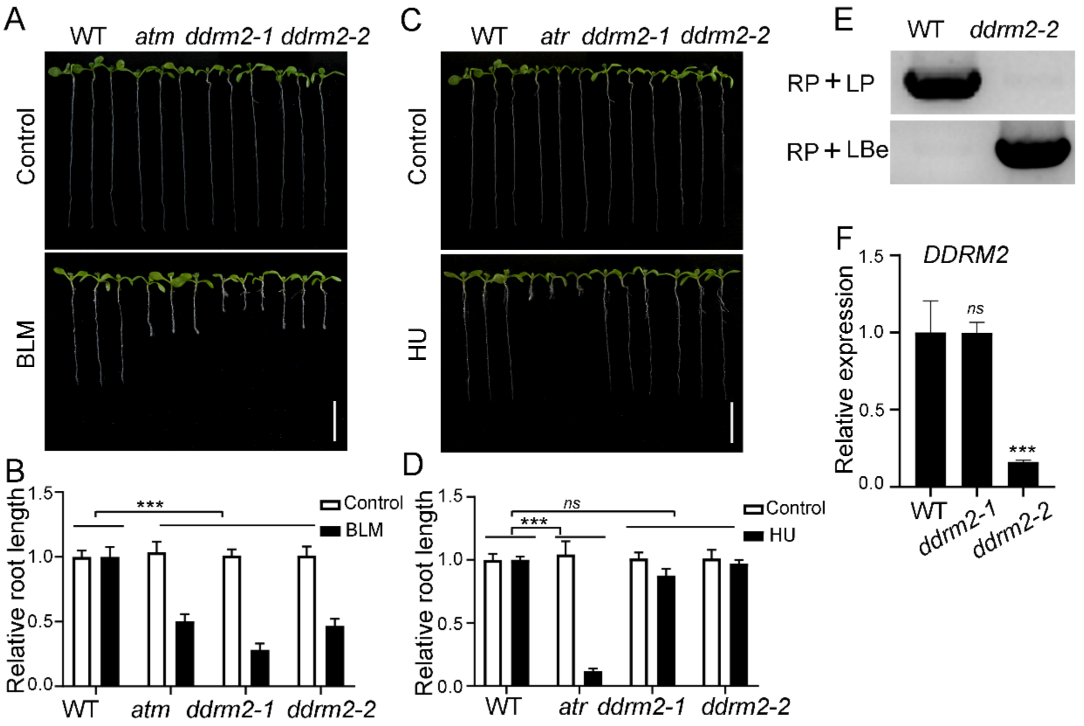


**Supplemental Figure S1.** The *ddrm2* mutants are hypersensitive to BLM, but not HU. A and C, Pictures of Arabidopsis seedlings grown on 1/2 MS with or without BLM (2.5 μM), HU (0.75 mM) for 8 days. Scale bar, 1 cm. B and D, The relative root length of the indicated plants. The relative root length data were represented as means ± SD (n = 10) relative to the values obtained under the control condition. The statistical significance was determined using two-way ANOVA analysis. ***, $P < 0.001$, *ns*, no significance. E, Genotyping results of *ddrm2-2*. LP and RP are primers flanking the insertion sites. LBe is the primer on the left border of T-DNA. F, The relative expression of *DDRM2* in *ddrm2* mutants analyzed by qRT-PCR. The *UBQ5* was used as a reference gene. Data represent mean ± SD (n = 3). All experiments were repeated three times with similar results.

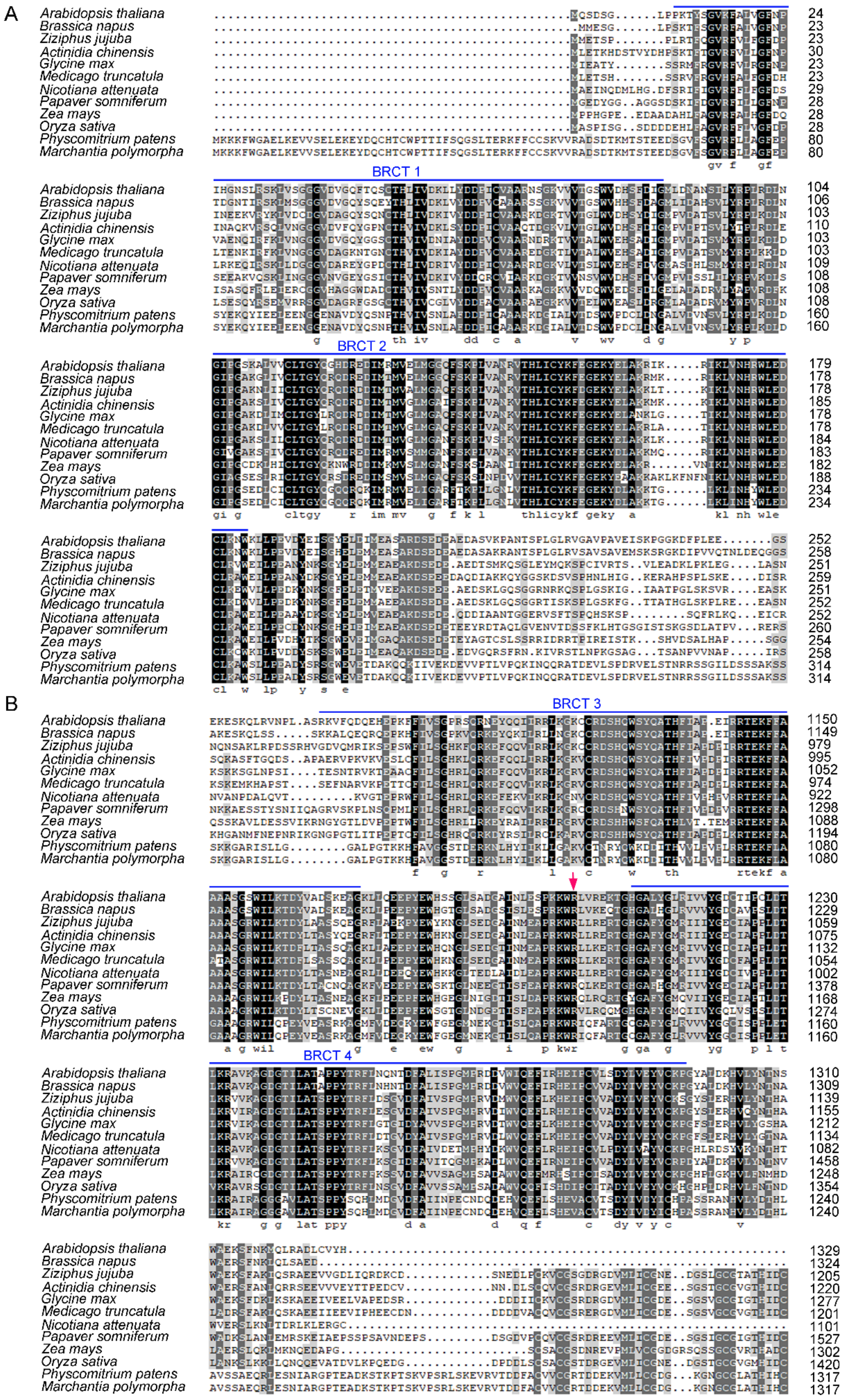
A
B
BRCT 1
BRCT 2
BRCT 3
BRCT 4
Arabidopsis thaliana
Brassica napus
Ziziphus jujuba
Actinidia chinensis
Glycine max
Medicago truncatula
Nicotiana attenuata
Papaver somniferum
Zea mays
Oryza sativa
Physcomitrium patens
Marchantia polymorpha

**Supplemental Figure S2.** The sequence alignment of DDRM2 and its orthologs in other plant species. A, N-terminus. B, C-terminus. Four BRCT domains were marked with blue lines. The mutated amino acid (Arginine 1201) in DDRM2 was marked with a red arrow. The alignment was conducted by CLUSTAL W.

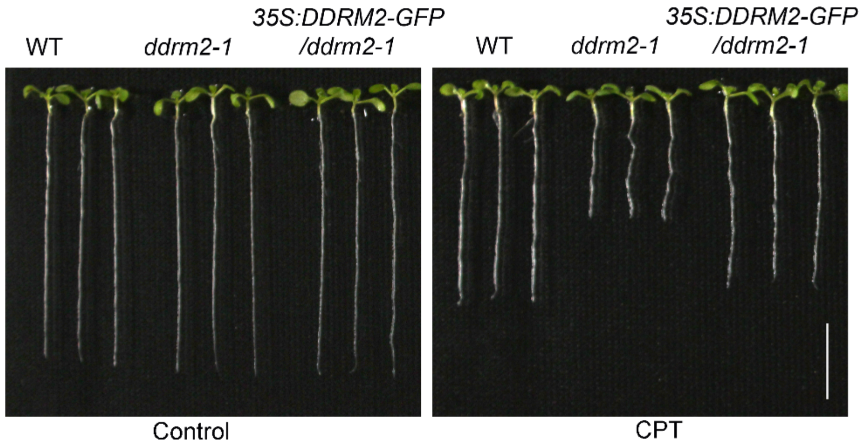


**Supplemental Figure S3.** DDRM2-GFP fusion protein driven by the *CaMV 35S* promoter can rescue *ddrm2-1*. These plants were grown on 1/2 MS media with or without CPT (15 nM) for 8 days. Scale bar, 1 cm.

**Supplemental Table S1 Candidate genes revealed by SIMPLE analysis**

| Chr | Position | Ref | Alt | Mutation_effect | Gene | AGI | CDS _change | Protein _change | Mut _ref | Mut _alt | WT _ref | WT _alt |
|---|---|---|---|---|---|---|---|---|---|---|---|---|
| 4 | 939631 | C | T | missense_variant | AT4G02110 | AT4G02110 | 3601C>T | Arg1201Trp | 0 | 21 | 54 | 0 |
| 4 | 1080504 | C | T | missense_variant | PMS1 | AT4G02460 | 7G>A | Gly3Arg | 0 | 25 | 81 | 0 |
| 4 | 1250291 | C | T | missense_variant | AT4G02800 | AT4G02800 | 74C>T | Ala25Val | 0 | 26 | 21 | 0 |
| 4 | 1311504 | C | T | missense_variant | AT4G02950 | AT4G02950 | 307C>T | Leu103Phe | 1 | 30 | 57 | 0 |

Chr, the chromosome number; Pos, the position of the SNP in each chromosome; Ref, the reference nucleotide in the reference genome; Alt, the alternative nucleotide in the mutant genome; mut_ref, the read number containing the reference nucleotide in mutant; mut_alt, the read number containing the alterative nucleotide in mutant; WT_ref, the read number containing the reference nucleotide in WT; WT_alt, the read number containing the alterative nucleotide in WT.

**Supplemental Table S2 Primers used in this study**

| Primer | Sequence | Purpose |
|---|---|---|
| atm-LP | ATCCATGTGGTTCAGTCTTGC | Genotyping |
| atm-RP | TTGGTATCCTGCAGAGGAAAG | Genotyping |
| LBe | GGAACAACACTCAACCCTATCTCG | Genotyping for Salk line |
| ddrm2-2-LP | GATGGTCTTTCTCTTCTGGGG | Genotyping |
| ddrm2-2-RP | CGCCAGAGACTGATACTTTGG | Genotyping |
| DDRM2-P07-F | ggatcctctagagtcgacctATGCAATCGGATTCGGG | Cloning |
| DDRM2-P07-R | caggtcgactctagaggatccATGGTACACACACAAATCTGCT | Cloning |
| ProDDRM2-GUS-F | ggggacaagtttgtacaaaaaagcaggcttcTTGTTGCCTGTTTGGTCC | Cloning |
| ProDDRM2-GUS-R | ggggaccactttgtacaagaaagctgggtcAAACGTAAATTGGGGATTG | Cloning |
| DDRM2-qPCR-F | GAGTTTATACGCCACGAAATCC | qPCR |
| DDRM2-qPCR-R | AATGGTACACACACAAATCTGC | qPCR |
| UBQ5-qPCR-F | GAAGATCCAAGACAAGGAAGGA | qPCR |
| UBQ5-qPCR-R | CTTCTTCCTCTTCTTAGCACCA | qPCR |
| ProDDRM2-0800-F | actatagggcgaattgggtacAGATTGAAAATTGTCTCGTG | Cloning |
| ProDDRM2-0800-R | ggccgctctagaactagtgTTTTTTTTTTGAAAAATTAGGGTTTTAT | Cloning |
| SOG1-2306-F | gagaacacgggggacgagctcggtaccATGGCTGGGCGATCATGGCT | Cloning |
| SOG1-2306-R | tctgatccaagggcgaattggtcgacTCAATCAGTCTTTCCAGTCCC | Cloning |
| LBx | ATATTGACCATCATACTCATTGC | Genotyping for GABI line |
| sog1-2-LP | AAAGTCCACCAAAGTGTGTCG | Genotyping |
| sog1-2-RP | GCCGATGCACTCATAATCTTC | Genotyping |
| IU8 gFW | ACTTATTTGCGTGTCTGTTACTT | Genotyping |
| IU8 gRV | GAAATCGATTACTAGCCACCACTC | Genotyping |
| IU8-T-DNA | TTGGACGTGAATGTAGACAC | Genotyping |

| | | |
|---|---|---|
| 2S-I-SceI#8 gFW | GGCGATCTCGACTCTCCTCTG | Genotyping |
| 2S-I-SceI#8 gRV | TCATCTCTTCCCGTAAAACCTCAA | Genotyping |
